# Genome-wide co-essentiality analysis in *Mycobacterium tuberculosis* reveals an itaconate defense enzyme module

**DOI:** 10.1101/2022.09.27.509804

**Authors:** Adrian Jinich, Sakila Z. Nazia, Andrea V. Tellez, Amy M. Wu, Ricardo Almada-Monter, Clare M. Smith, Kyu Rhee

## Abstract

Genome-wide random mutagenesis screens using transposon sequencing (TnSeq) have been a cornerstone of functional genetics in *Mycobacterium tuberculosis* (*Mtb*), helping to define gene essentiality across a wide range of experimental conditions. Here, we harness a recently compiled TnSeq database to identify pairwise correlations of gene essentiality profiles (i.e. co-essentiality analysis) across the *Mtb* genome and reveal clusters of genes with similar function. We describe selected modules identified by our pipeline, review the literature supporting their associations, and propose hypotheses about novel associations. We focus on a cluster of seven enzymes for experimental validation, characterizing it as an enzymatic arsenal that helps *Mtb* counter the toxic effects of itaconate, a host-derived antibacterial compound. We extend the use of these correlations to enable prediction of protein complexes by designing a virtual screen that ranks potentially interacting heterodimers from co-essential protein pairs. We envision co-essentiality analysis will help accelerate gene functional discovery in this important human pathogen.

## Introduction

*Mycobacterium tuberculosis* is the deadliest human bacterial pathogen, yet the function of the vast majority of its genes remains unknown (Fig. 1). Statistical grouping of genes based on conditional essentiality or fitness datasets helps guide functional annotation ^1–6^, as clustering genes into modules effectively reduces the dimension of the annotation challenge. Such co-essentiality (or co-fitness) analysis, which generates genes clusters or modules based on correlations in conditional fitness profiles across large numbers of standardized screens, has been applied to a wide variety of organisms. In microbiology, co-essentiality analysis of many bacterial species has burgeoned with the development of randomly bar-coded transposon sequencing (RBTnSeq) ^7,8^. In human and cancer cell lines, pooling CRISPR-based fitness data from hundreds of genome-wide essentiality screens has revealed co-essential gene modules and helped accelerate gene functional discovery ^1,4,9^.

**Figure 1:**
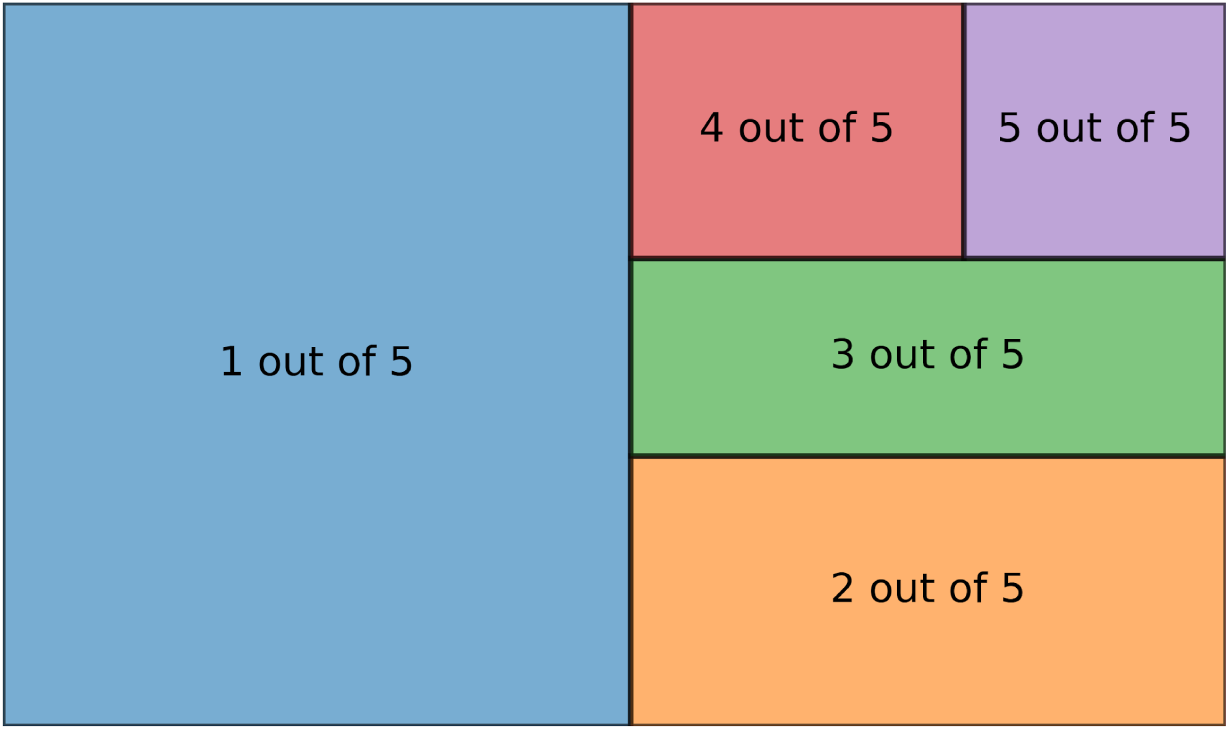
The state of protein functional annotation of the *Mycobacterium tuberculosis (Mtb)* proteome according to UniProtKB. The full rectangle represents the *Mtb* proteome, and each colored sub-section are the fractions with different UniProtKB annotation scores, a heuristic measure of the annotation content of a protein entry. These range on a scale from 1 to 5, with protein entries with experimental evidence scoring higher than annotations inferred from homology ^74^.

Despite the established trajectory of transposon sequencing (TnSeq) in *Mtb* functional genetics ^10–13^, co-essentiality analysis has yet to be systematically applied for *Mtb* functional genomics. This is partly due to the lack of a resource that makes *Mtb* TnSeq conditional essentiality data standardized and easily accessible. We recently compiled a database of *Mtb* TnSeq screens across dozens of experimental conditions ^14^. Here we harness our Mtb TnSeq screen database and a recent statistical method for co-fitness analysis based on generalized least squares (GLS) ^9^ to systematically search for TnSeq conditional fitness correlations that hint at gene function and modular associations. This reveals both known and novel complexes and metabolic functional modules. We focus on a cluster of 7 enzymes for experimental validation, and demonstrate that they are part of an enzymatic arsenal to defend against the host’s attack with itaconate, a key antibacterial compound. Finally, we extend the use of these co-fitness associations and design a virtual screen that takes as input gene pairs with strongly correlated co-essentiality profiles and, using AlphaFold2-multimer ^15^ and a recent predicted protein complex structure quality scoring function ^16^, predicts and ranks them according to the likelihood that they form protein-protein heterodimer complexes. We envision co-essentiality analysis will help future efforts to decipher gene function in *Mtb*. To support and enhance these efforts, the co-essentiality analysis is now accessible as part of our *Mycobacterium tuberculosis* transposon sequencing database (MtbTnDB) online portal, providing wide access to interactive exploration of gene correlations.

## Results

### Co-essentiality analysis of Mtb TnSeq profiles confirms well-established metabolic pathways and reveals functional nuances

Identifying a gene’s conditional essentiality profile, or the conditions where a gene is essential, can help elucidate its function. Genes with similar conditional essentiality profiles are likely co-regulated and work together in a functional module. Using our TnSeq database, MtbTnDb, we generated conditional essentiality profiles, defined as the set of conditional essentiality calls across all standardized TnSeq screens (i.e. a 125-dimensional vector), for each gene in the Mtb genome ^14^. We discerned pairs of genes with similar TnSeq conditional essentiality profiles by identifying pairwise comparisons that result in significant generalized least squares (GLS) regression coefficients and strong linear relationships ^9^. Clustering significant gene pairs with overlapping gene pairs revealed several functional modules, including clusters of genes that are known to interact with each other, such as the phthiocerol dimycocerosate (PDIM) operon (Fig. 1). This confirmed that GLS regression analysis can detect functional gene relationships. We also found clusters of genes that included well-characterized genes with uncharacterized genes or other characterized genes with no apparent interactions. This highlights the potential of using our TnSeq database and GLS regression analysis to identify novel protein interactions and generate novel hypotheses about the function of orphan proteins or alternative functions for well-characterized proteins.

Here, we describe selected functional modules identified by our GLS pipeline and review the literature to support their associations and propose hypotheses about novel associations (Fig. 2).

**Figure 2:**
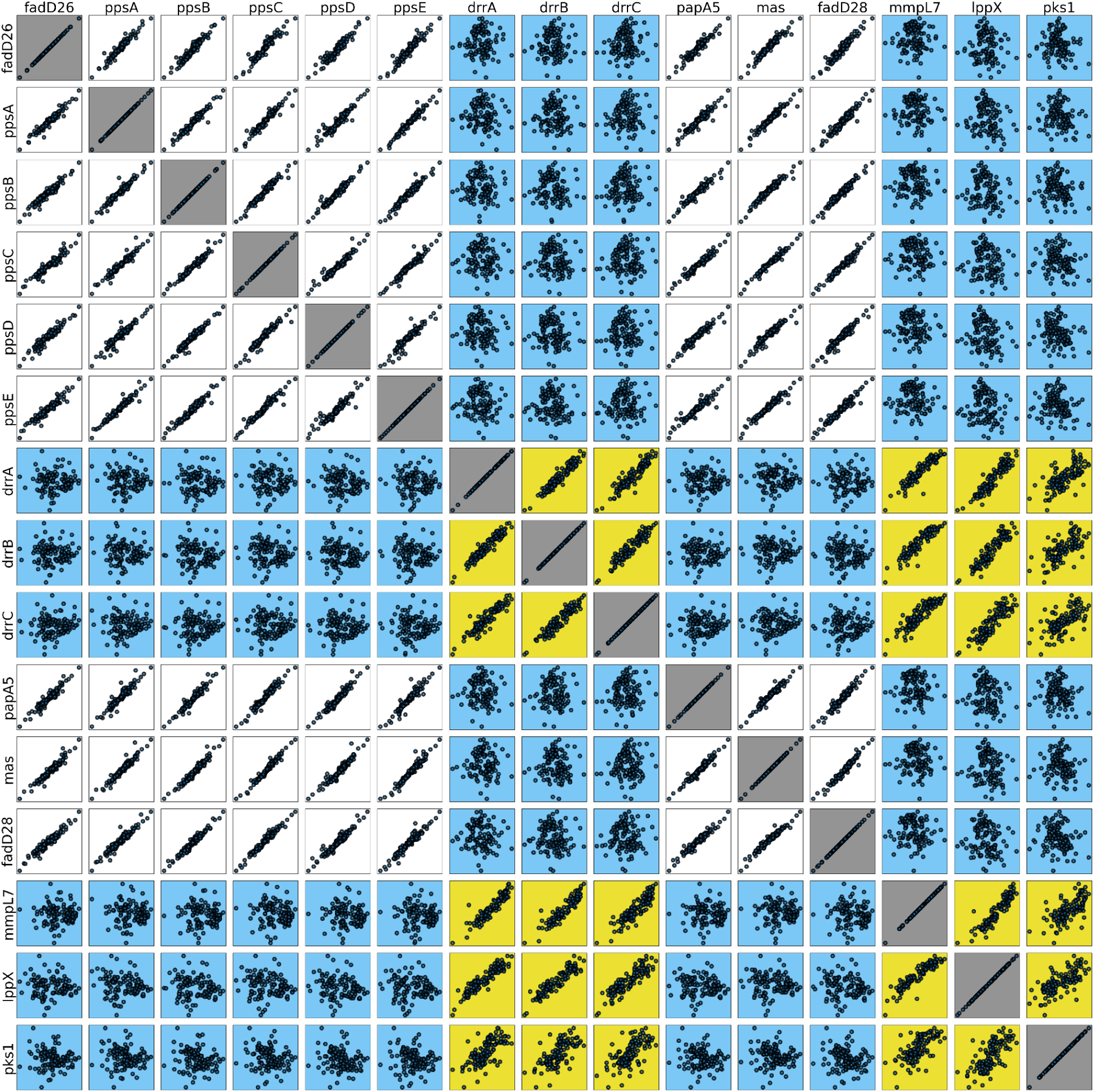
Patterns of co-essentiality in *Mtb* PDIM metabolism and transport. Pairwise correlations (or lack of) in conditional essentiality profiles between PDIM biosynthesis genes (white); between PDIM transport genes (yellow); and between PDIM transport and biosynthesis genes (blue). The x- and y-axes are in log(2) fold-changes in transposon insertion counts. Each dot in each scatter plot corresponds to a single TnSeq screen, and is obtained from comparing counts in an experimental condition relative to a control condition as standardized in the MtbTnDB.

#### PDIM Synthesis and Transport

The most notably defined module identified by our GLS pipeline included genes of the PDIM operon, responsible for the biosynthesis of the virulence lipid, PDIM. This module consists of genes involved in the synthesis of PDIM phthlidone moiety, (FadD26, ppsA-E/Rv2930-Rv2935) the synthesis of mycoseric acid, (FadD28, mas/Rv2940c-Rv2941c), and the esterification of two mycoseric acids and one pthiolodone to form PDIM (PapA5/Rv2939) ^17,18^ (Fig. 2, Module 1a). Interestingly while the co-essentiality profiles of PDIM biosynthesis genes correlate in a pairwise manner, they do not correlate with profiles of PDIM transport genes, which are located adjacently to PDIM biosynthesis genes in the *Mtb* genome. Instead, PDIM transport genes, including daunorubicin-dim-transport resistance ABC transporter locus (*drrA-C/*Rv2936-8), Mmpl7 transporter (Rv2942), and the lipoprotein LppX (Rv2945c) appear as a separate module from PDIM biosynthesis genes (Fig 2, Module 1b). Mmpl7 and drrA-C have been shown to be important in PDIM localization to the cellular envelope, and even suggested to form a complex with PDIM biosynthesis genes ^19–21^. Likewise, lppX is also proposed to facilitate PDIM localization by binding and transporting the lipid across the plasma membrane to the outer leaflet ^22^.

While these genes’ importance in PDIM transport has been thoroughly investigated, their role in transport of other lipids is unclear ^23^. We speculate that while Mmpl7, drrA-C, and lppX are involved in PDIM transport, they could predominantly function to transport other lipids, such as phenolic-glycolipids (PGL). This is supported by studies showing that genetic knockout of Mmpl7 prevents PGL localization to cell membrane, as well as, our GLS pipeline clustering PDIM transport genes with pks1 (*Rv2946c*), shown to be essential in PGL synthesis ^23–25^. Thus, our GLS pipeline confirms previous literature describing PDIM biosynthesis, but offers new insight on alternative roles of transport proteins that are currently only annotated to be involved in PDIM metabolism.

#### Mycobactin Synthesis

In addition to PDIM biosynthesis, another module identified by the GLS pipeline involves genes of the mycobactin siderophore (MBT) biosynthetic pathway. MBT biosynthesis genes are located in two separate operons, mbt-1 and mbt-2. While the genes in the mbt-1 operon (mbtA-I/Rv2378c-86c) use polyketide synthase/nonribosomal peptide synthetase chemistry to build the heterocyclic scaffold of mycobactin, the genes in the mbt-2 operon (mbtK-N) are used to modify the siderophore with aliphatic chains to form mycobactin derivatives ^26,27^.

We find that the TnSeq profiles of all proteins in the mbt-1 operon, except mbtH, are tightly correlated (Fig. 3, Module 2, Supp. Fig. 1a). The lack of a significant correlation with mbtH is likely due to its small size (71 residues) which likely results in noisy TnSeq insertion counts. Strikingly, the TnSeq profiles of the mbt-1 genes do not correlate with those of mbt-2 genes, and the mbt-2 genes’ TnSeq profiles do not correlate amongst themselves. We speculate that genes of the mbt2 operon have physiologic roles beyond synthesis of mycobactin derivatives. This is consistent with previous literature reporting that mbtK, but not mbtN, is essential for growth in iron limitation and also influences the composition of the global phospholipid landscape ^28^. Similar to mbt-2 genes, the TnSeq profile of irtAB, located adjacent to the mbt-2 locus annotated as mycobactin transporter also does not correlate with profiles of mbt-1 and mbt-2 genes ^29^, suggesting an alternative physiological function in addition to mycobactin transport. Thus, our GLS pipeline confirmed previous literature describing the mycobactin scaffold, while revealing nuances about mycobactin modification and transport.

**Figure 3:**
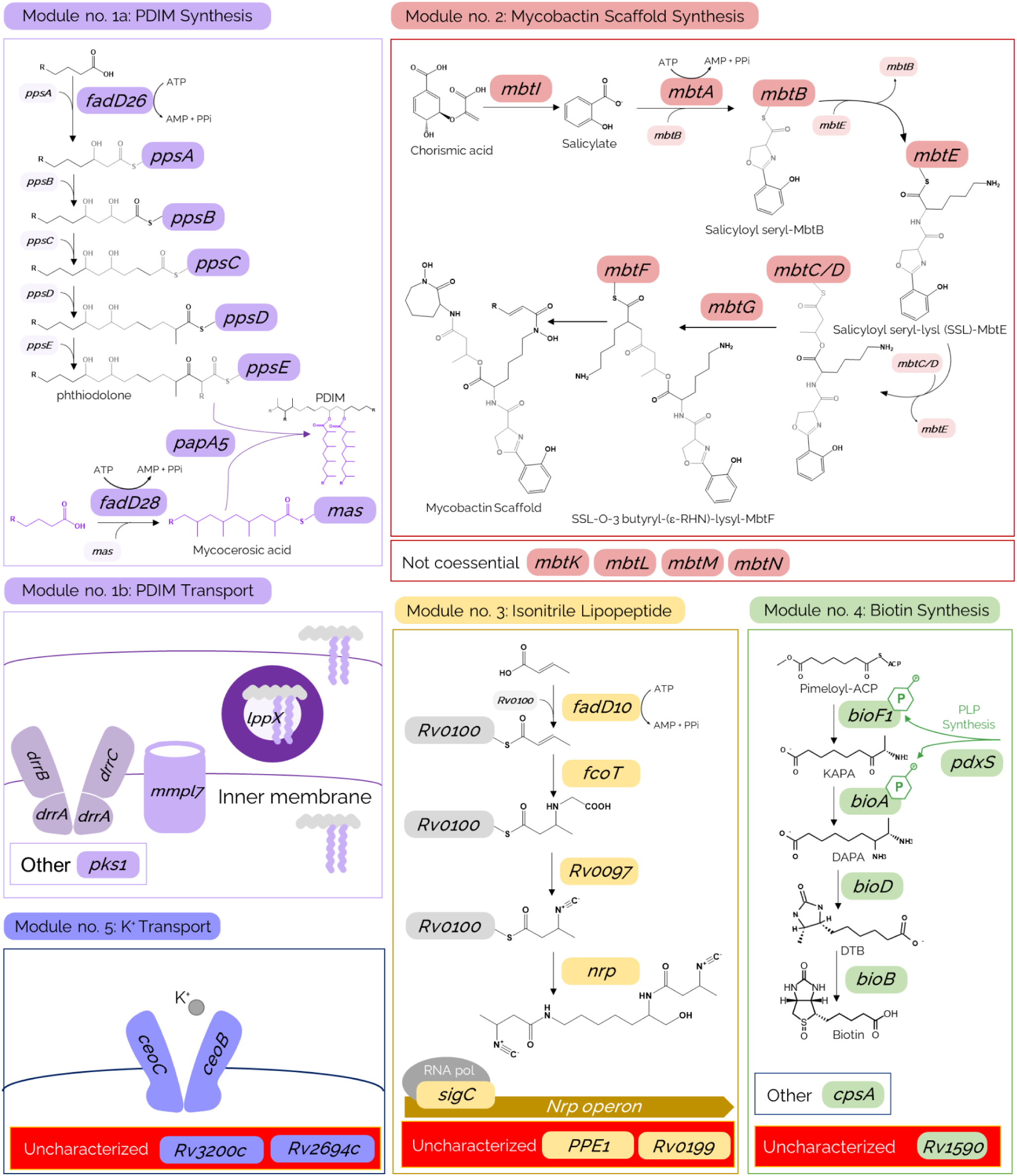
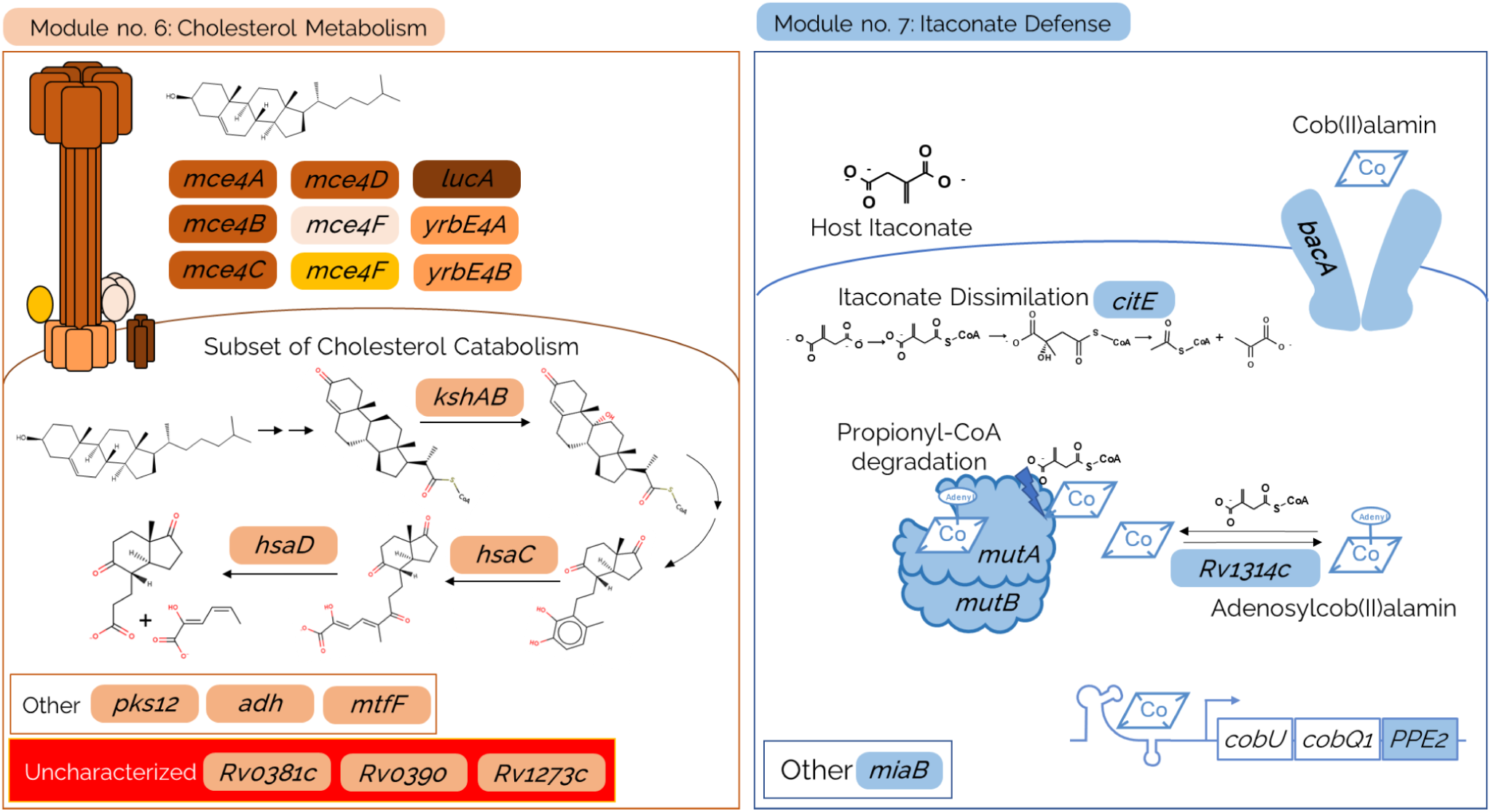
Gene functional modules identified by co-essentiality patterns in TnSeq data. Illustration of select modules, including (1a) PDIM synthesis, (1b) PDIM transport, (2) mycobactin scaffold synthesis, (3) Isonitrile lipopeptide synthesis, (4) biotin synthesis, (5) potassium (K^+^) transport, (6) cholesterol metabolism, and (7) itaconate defense.

#### Isonitrile Lipopeptide Synthesis

Another module identified by the GLS pipeline (Fig. 3, Module 3, Supp. Fig. 1b) consists of genes in the nonribosomal peptide synthase (nrp) operon, genes Rv0096- Rv0101, except Rv0100, as well as, the alternative sigma factor sigCC (Rv2069). We speculate that the TnSeq profile of Rv0100 was not correlated with the rest of the nrp operon because of its small size (78 residues) and statistical potential for noisy transposon insertion count data. The Rv0096-Rv0101 operon codes for enzymes that synthesize an isonitrile lipopeptide, which has a putative role in biofilm development and copper acquisition during copper limitation ^30–32^. Recent work implicates PPE1 as a functional part of the cluster, supporting its co-essentiality with the rest of the genes observed in our analysis ^32^. SigC has been implicated in the transcriptional activation of the nrp operon in response to transition metal limitation based on studies of SigC overexpression and DNA binding, and an observed growth defect of an *Mtb* strain with sigC knocked out in copper limited media ^33–35^. Thus, this module demonstrates how co-essentiality captures regulatory interactions without relying on canonical gene expression profiling.

#### Biotin Synthesis

Our GLS pipeline also identified a module of genes that includes a subset of biotin biosynthesis genes, including bioA, bioF1, bioD (Rv1568-70) and bioB, as well as, pyridoxal 5’-phosphate (PLP) synthase (PdxS/Rv2606c), cpsA (Rv3484), and Rv1590, an uncharacterized protein (Fig. 3, Module 4, Supp. Fig. 1c). Biotin synthesis can be divided into two stages: (1) pimeloyl-ACP synthesis and (2) its assemblage into biotin’s bicyclic rings ^36^. Our GLS pipeline only identifies co-essentiality between genes from the second stage of biotin synthesis and PdxS, involved in PLP synthesis, a cofactor for bioF1 and bioA ^37^.

The exclusion of putative genes of the first stage of biotin biosynthesis suggests that pimeloyl-ACP synthesis is independent of biotin assembly. Pimeloyl-ACP synthesis is still poorly understood in *Mtb* but is suggested to be mediated by Tam (Rv0294), an O-methyltransferase homologous to the *Escherichia coli* gene BioC, part of the BioC-BioH pimelate synthesis pathway in *E. coli* ^38,39^. However, unlike other biotin synthesis genes, Tam was not found to be conditionally essential in any TnSeq screens used in our GLS comparison. We speculate that *Mtb* may encode redundant pathways for pimeolate synthesis that are found in other organisms ^36^. The significant co-fitness correlations observed between CpsA and biotin synthesis is unclear, as CpsA is involved in linking arabinogalactan to peptidoglycan as well as fatty acid import ^40–42^.

#### Other Modules

In addition to the four modules described above, our GLS pipeline identified a module including genes involved in cholesterol import and metabolism (Fig. 3, Module 6, Supp. Fig. 1e). It consists of genes part of the Mce4 operon (Rv3492c - Rv3500c), responsible for cholesterol import, a subset of cholesterol catabolism genes, kshA (Rv3526) and hsaC- hsaD (Rv3658c - Rv3659c), and genes involved in alcohol metabolism, adhB (Rv0761c), an alcohol dehydrogenase, and mftF (Rv0696), pre-mycofactocin glycosyltransferase ^43–45^. Alcohol metabolism genes are likely co-essential with cholesterol metabolism genes, since conditions with cholesterol supplemented growth media also contain alcohol ^46^. Other genes part of this module include pks12 (Rv2048c), which synthesizes beta-D-mannosyl phosphomycoketide, and three uncharacterized proteins Rv0381c, Rv0390, and Rv1273c ^47^.

Our GLS pipeline also identified a module involving genes in the Trk operon, ceoB (Rv2691) and ceoC (Rv2692), (Fig. 3, Module 5, Supp. Fig. 1d) which mediate potassium transport, as well as two uncharacterized proteins, *Rv2694c* and *Rv3200c* ^48^. The modules we describe here are only a select few of the many associations our co-essentiality approach identifies. These examples demonstrate the wide array of potential hypotheses our GLS pipeline can generate.

### Co-essentiality in Mtb TnSeq profiles reveals an enzymatic arsenal to defend against host itaconate

To test the robustness of the modules identified by our co-essentiality analysis, we selected one module for further experimental validation. We focused on a module that includes a genes involved in cobalamin (B12)-dependent metabolism, including bacA (Rv1819c), a cobalamin transporter ^49–51^, ATR (Rv1314c) corrinoid adenosyltransferase, and mutA-B (Rv1492-3) the cobalamin-dependent enzyme, methylmalonyl CoA mutase (MCM). This module also contains the following genes: citE (Rv2498c), a β-hydroxyacyl-CoA lyase (β-hCL) ^52^, PPE2 (Rv0256c), and miaB (Rv2733c) (Fig 2, Module 7, Supp. Fig. 1e).

Speculating on why these genes form a co-essential cluster, we recognized that they are related to MCM and its perturbations during itaconate toxicity, a host immunometabolite released by macrophages ^53^. MCM activity, which involves the isomerization of methylmalonyl-CoA to succinyl-CoA during propionate catabolism, relies on the cofactor, adenosylcobalamin (AdoCbl). During its catalytic mechanism, MCM homolytically cleaves AdoCbl to cob(II)alamin and Ado**·.** ATR acts as a chaperone for MCM by off-loading and stabilizing the radical intermediates and then directly adenylating cob(II)alamin to AdoCbl in MCM’s active site ^54,55^. Itaconate poisons the MCM system through its CoA derivative, itaconate-CoA (I-CoA), by stabilizing the Ado**·** radical through formation of a biradical dAdo-I-CoA adduct in MCM’s active site ^53^. This prevents AdoCbl regeneration and renders MCM inactive, preventing *Mtb* from metabolizing propionate, a major carbon source for *Mtb* in the host environment.

We hypothesized that *Mtb* has evolved a response to itaconate stress which employs genes identified by our module, including ATR, citE, PPE2, and miaB. ATR has been shown to repair MCM after I-CoA inactivation in *in vitro* studies of MCM catalysis co-incubated with I-CoA and ATR ^53^. CitE, which encodes a β-hCL also contributes to itaconate defense by catabolizing I-CoA into acetyl-CoA and pyruvate ^52^.

The roles of PPE2 and miaB in itaconate defense are less clear. PPE2 expression is co-regulated with cobalamin synthesis genes, cobQ1 and cobU by a cob(II)alamin-sensitive riboswitch upstream of the PPE2-cobQ1-cobU operon ^49^. Although PPE2 is regulated by cob(II)alamin, its role in cob(II)alamin metabolism has yet to be elucidated. PPE2 is annotated to be a secreted protein that contains a nuclear localization signal and DNA-binding motif that allows it to downregulate transcription of immunomodulatory genes in macrophages, including nitric oxide (NO) and reactive oxygen species (ROS) synthesis genes ^56,57^. PPE2 specifically binds to the GATA-1 sequence, also found in the promoter region of immune responsive gene (Irg1) which encodes aconitase decarboxylase, responsible for itaconate production in host immune cells. One possibility is that PPE2 responds to itaconate-mediated AdoCbl depletion by downregulating aconitase and itaconate production in host macrophages. MiaB is less well characterized and is annotated as a putative tRNA methylthiotransferase based on sequence homology and the presence of a radical S-adenosyl-L-methionine (rSAM) domain. Since rSAMs have been shown to thiomethylate small molecules as well as tRNAs, one possibility is that miaB responds to itaconate toxicity by thiomethylating itaconate or I-CoA, preventing dAdo-I-CoA adduct formation in the MCM active site ^58^.

To determine if this functional module represents a mechanism of *Mtb’*s defense against itaconate toxicity, we tested whether ATC-induced CRISPRi silencing of these genes exacerbated the growth attenuation caused by itaconate when *Mtb* is grown *in vitro* with propionate as the sole carbon source ^53^. Importantly, this experimental condition is not included in the underlying *Mtb* TnSeq database used to generate the co-essential clusters, demonstrating how our analysis leads to hypotheses involving unexplored experimental setups. In the absence of gene knockdown (-ATC), all strains experienced growth defects in 10 *μ*M itaconate, while growth in minimally affected in 1 *μ*M itaconate (Fig 4a, b). However, the genetic knockdown strains, *mutB-KD, mutA-KD, citE-KD, rv1314c-KD, ppe2-KD,* and *bacA-KD* experienced an additional growth defect in 1 *μ*M itaconate (Fig 4c). Knockdown of some genes also caused growth defects in glycerol media, however, to a significantly lesser extent to growth defects in 1 *μ*M itaconate (Fig 4c,d). This suggests these genes are important specifically for itaconate defense during MCM-mediated growth, rather than for growth in general. Notably, the growth defect of the *ppe2-KD* suggests that PPE2 contributes to itaconate defense independent of its possible role in regulating host immunomodulatory genes. The uniform response of these genes to downregulation in the presence of itaconate corroborates our GLS pipeline’s clustering of these genes and data demonstrate the utility of the co-essentiality approach in identifying novel functions for well-characterized and uncharacterized proteins.

**Figure 4:**
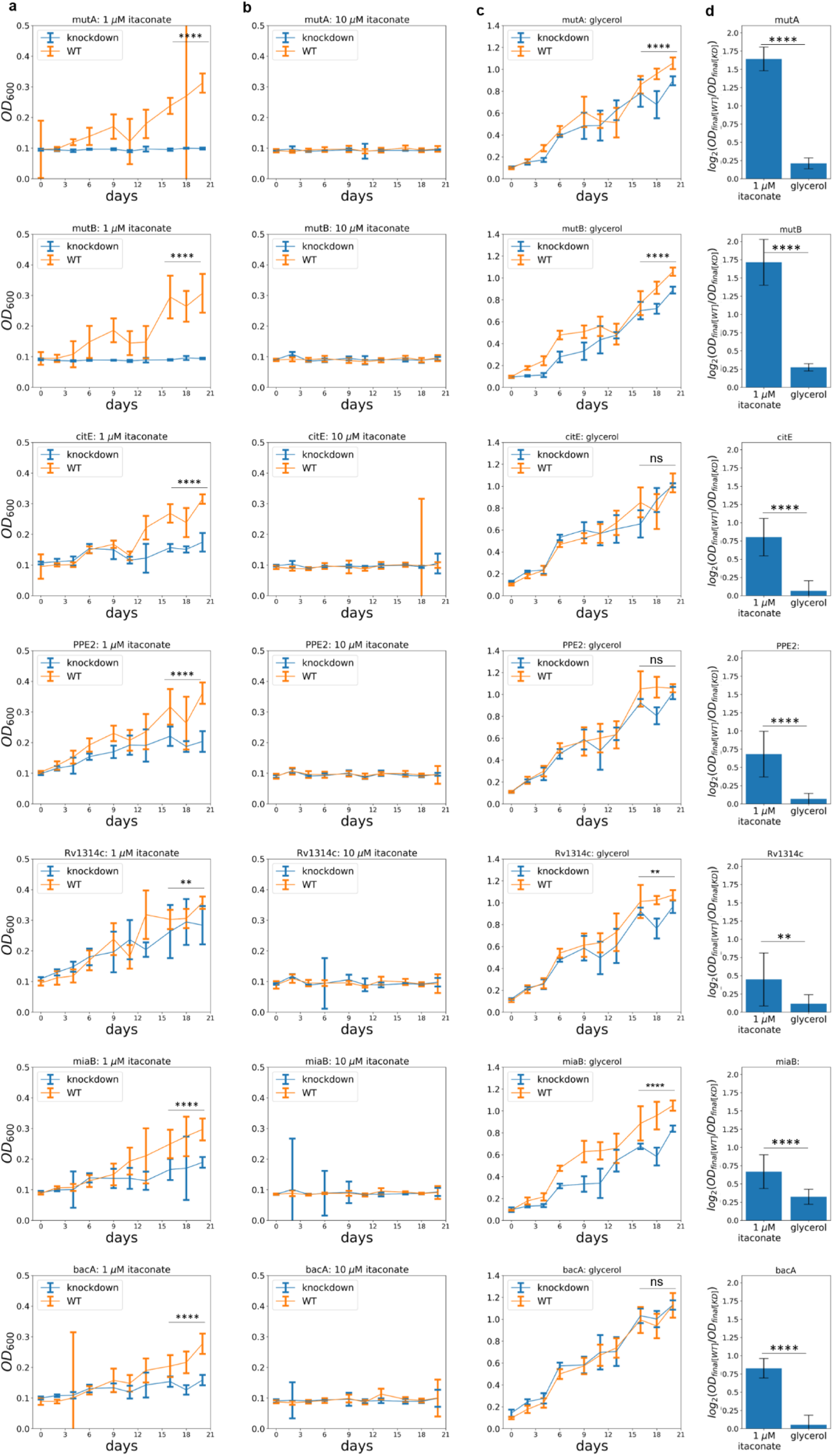
Knockdown of genes in itaconate defense module exacerbates itaconate mediated growth defect. Optical density at 600 nm (OD_600_) of *Mtb* strains with ATC-induced CRISPRi silencing system, *mutA KD, mutB KD, citE KD, Rv1314c KD, miaB KD,* and *bacA KD* liquid cultures grown with presence (blue) and absence of anhydrotetracycline (ATC) (orange) in (a) 0.02% propionate and (b) 1, (c) 10 *μ*M itaconate, or glycerol over 20 days (N = 9, p < 0.05, student’s t-test, Error bars = SD). (d) Relative fitness defect of gene KD as measured by log_2_ ratio of the day 21 OD_600_ of the strains grown with ATC to OD_600_ of the strains grown without ATC in 1 *μ*M itaconate and glycerol (N = 9, p < 0.05, paired student’s t-test, Error bars = SD)

Next we sought to confirm whether the observed co-essentiality of genes in the itaconate-defense module is relevant in *in vivo* mouse infection models. We analyzed TnSeq fitness profiles for these 7 genes using only a subset of screens involving infection of mouse strains of the collaborative cross (CC) panel. Analysis of TnSeq data for infected CC-panel mice has shown their genetic diversity is representative of the range of phenotypic responses to *Mtb* infection ^59,60^. When the search for significant co-fitness signals is restricted to only TnSeq screens from the *in vivo* CC-panel dataset, we find significant TnSeq profile correlations involving 6 of the 7 genes in the module (FDR-corrected p-values < 0.001, Table S3). The CC-panel mouse strains with the strongest fitness defects across genes in the module (< *log_2_FC* > ≤ − 1. 5) include (Fig. 5): CAST/EiJ, CC005/TauUnc, CC039/Unc, Cybb^-/-^, Nos2^-/-^, CC001/Unc, and CC031/GeniUnc. Interestingly, *Nos2*^−/−^ bone marrow-derived macrophages (BMDMs), have been shown to have higher itaconate levels than wild-type (WT) BMDMs ^61^. Future work could aim to identify whether the genetic determinants underlying these mouse strains’ response to infection (with *Mtb* strains bearing mutations in the identified genetic module) are related to itaconate production.

**Figure 5:**
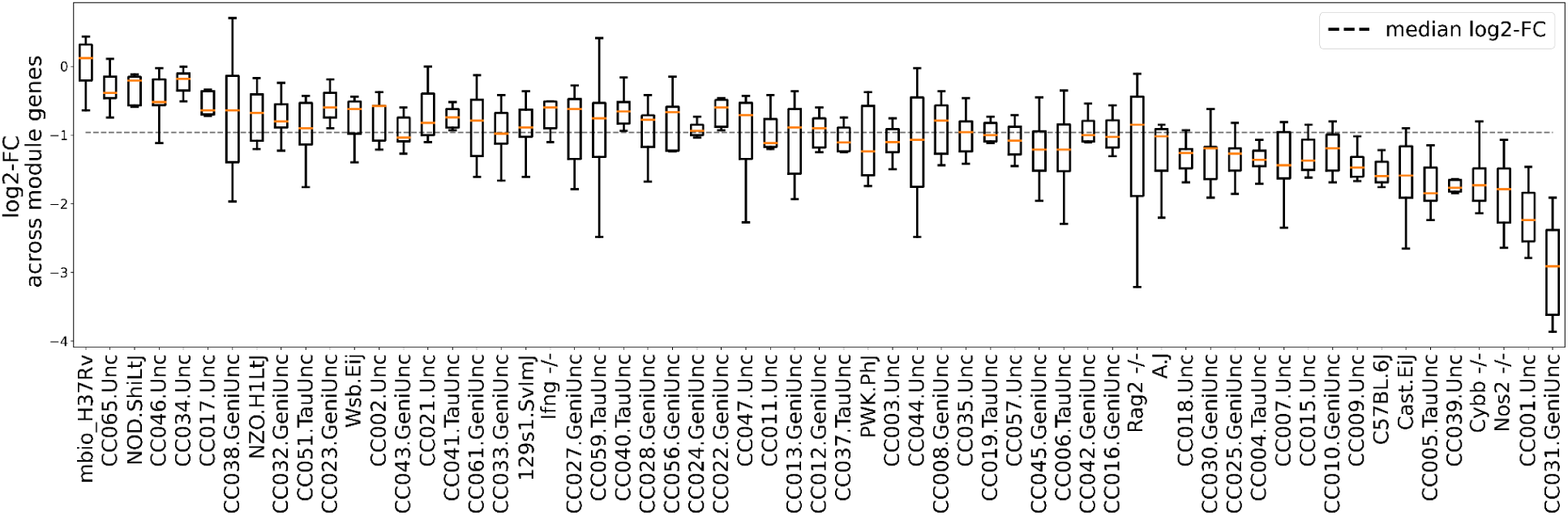
Genes in the itaconate defense module are co-essential in a subset of TnSeq screens involving collaborative cross (CC) mouse infection models. Box plot visualizing the distribution of log2 fold changes (log2-FC) of transposon insertion counts across genes belonging to the itaconate defense module (mutA, mutB, citE, ppe2, rv1314c, bacA, and miaB) in each TnSeq screen (mouse strain) belonging to the CC panel. The dashed line corresponds to the median log_2_ FC for the 7 itaconate defense module genes across all CC panel TnSeq screens.

### Computational prediction and scoring of protein heterodimer structures ranks interacting protein pair candidates

Motivated by the observation that some of the strongest GLS correlations observed reflect known physical protein-protein interactions (PPIs), we hypothesized that co-essentiality analysis could be used as an initial filter to computationally screen candidate pairwise PPIs. We reasoned this would prune the set of all possible protein pairs (≈8 million in the case of Mtb) to the subset with experimentally detectable correlated fitness defects. We therefore designed a virtual screen to detect and rank potential interacting protein pairs candidates obtained from GLS co-essentiality (Methods). Our virtual screen combines AlphaFold2 (AF2)-based complex structure prediction ^15,62^ with a recent protein complex scoring function, pDockQ ^16^. The pDockQ score, which has been shown to distinguish truly interacting from non-interacting proteins, is a parametrized logistic function of the product of two terms: the (logarithm of) the number of predicted interface contacts; and AF2’s plDDT quality values for amino acid residues at the interface. We reasoned that, while simulated complex structures may not yet consistently achieve the high accuracy of single protein structures, true protein-protein interactions might have, on average, significantly higher pDockQ scores than adequately controlled random pairs of proteins.

Towards this end, we predicted protein pair heterodimer structures using AF2-multimer (as implemented in ColabFold) for 130 protein pairs with significant GLS co-essentiality correlations. We scored the predicted heterodimer structures using pDockQ. To calibrate the pDockQ scores for PPI candidates against what would be expected by chance, we also ran AF2-multimer calculations for an equivalent number of random pairs of proteins with a similar size distribution. Fig. 6 shows the pDockQ scores obtained for both sets of protein pairs.

**Figure 6:**
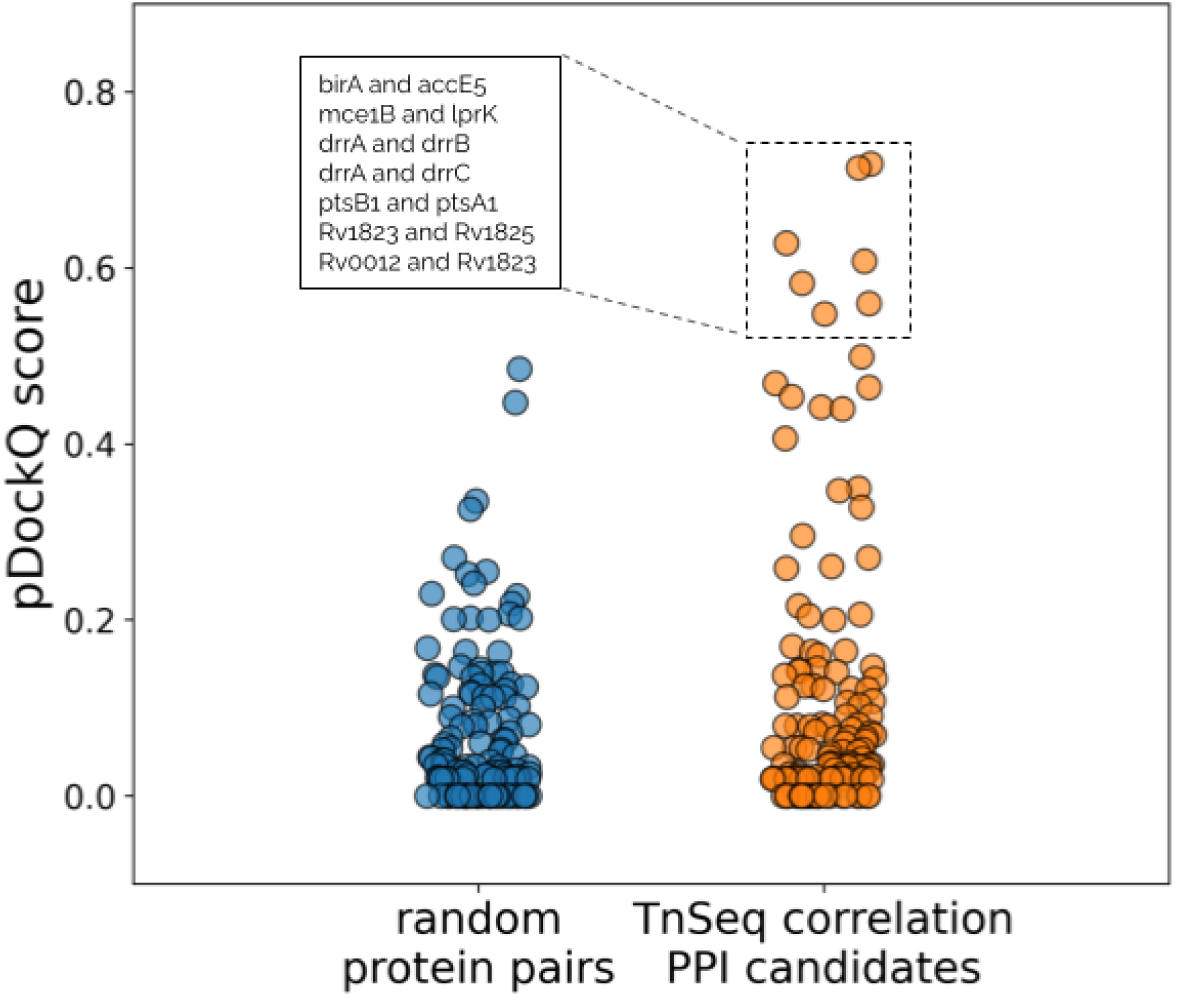
Distribution of pDockQ scores for AlphaFold2 heterodimer complex predictions from randomly selected protein pairs in comparison to protein pairs with strong conditional essentiality profile correlations. (Blue): pDockQ score distribution for randomly selected pairs of proteins; (Orange): pDockQ score distribution for PP-pairs with significant GLS co-essentiality correlations. Box highlights candidate protein pairs with the highest pDockQ scores.

We find 7 predicted heterodimer complexes that satisfy the strict threshold of having pDockQ scores higher than any pair of proteins from the randomized control set. Five of these heterodimers consist of proteins of known function. Two high-scoring heterodimers involve pairwise interactions between three PDIM transport genes of the annotated daunorubicin-dim-transport resistance ABC transporter locus: drrA (Rv2936) with drrB (Rv2937) (pDockQ=0.61), and drrA with drrC (Rv2938) (pDockQ=0.59). Two proteins, the ATP-binding protein ptsB1 (Rv0820) and the transmembrane-domain containing protein ptsA1 (Rv0930) (pDockQ=0.560), are annotated according to UniProt as *probably* part of the ABC transporter complex PstSACB involved in phosphate import ^63^. Another heterodimer with high pDockQ score involving well-annotated genes is birA (Rv3279c) with accE5 (Rv3281) (pDockQ=0.548). BirA catalyzes the transfer of biotin onto the biotin carboxyl carrier protein (BCCP) domain of acetyl-CoA carboxylase, converting it to active holo-BCCP ^64^, while accE5 is known to stimulate activity of the biotin-dependent acyl-CoA carboxylase complex ^65,66^. The remaining high-scoring protein pairs involve proteins of unknown function, including the highest scoring heterodimer is mce1B (Rv0170) with lprK (Rv0173) (pDockQ = 0.718), and the following two gene pairs: Rv1823 with Rv1825 (pDockQ=0.714) and Rv0012 with Rv1823 (pDockQ=0.628).

## Discussion

In this work, we identified gene co-essentiality relationships in *Mtb* by exploiting available TnSeq datasets and statistical methods that allow us to find significant pairwise correlations of gene TnSeq profiles. Clustering of co-essential genes reveals functional modules containing proteins with different degrees of functional annotation. Well-characterized proteins in a gene cluster can be used to generate hypotheses about orphan enzymes in the same cluster. We propose hypotheses regarding several identified clusters, and experimentally validated the co-essentiality of a cluster in growth media containing propionate and itaconate, despite the fact that this experimental condition was not part of the TnSeq database that revealed the correlations.. We show that these genes are co-essential when propionate, which drives mutAB metabolism, is the main carbon in the presence of itaconate. Co-essentiality is not observed in standard growth media, and hence, this seven gene module is likely involved in *Mtb*’s response to itaconate stress. Combining our approach with biochemical methods of enzyme discovery, such as activity based metabolic profiling (ABMP) can further confirm this physiological role ^67,68^.

We couple our co-essentiality approach with a virtual screen for heterodimer protein-protein complex detection. This screen applies a combination of AlphaFold2 structure prediction with the pDockQ scoring function to our co-essential gene pairs and shows that co-essential gene pairs are enriched for statistically significantly high complex quality scores. We showed that co-essentiality coupled with PPI predictions can successfully identify known complexes as well as predict novel interactions. The predicted interaction between the highest-ranked heterodimer complexes could be validated experimentally using pull-down assays or other PPI detection approaches. While our co-essentiality approach identified several strong PPI candidates, our computational PPI approach is limited to heterodimers, when many heterodimers are in reality part of higher-order complexes. Even so, our approach identified several heterodimers part of larger complexes (e.g. drrA, drrB, and drrC), highlighting its role in protein complex prediction beyond protein pairs. To more effectively capture larger, more intricate complexing interactions, we envision combinatorially exploring different stoichiometries of complex-forming candidates, although at a higher computational cost. Finally, we note that co-essentiality does not necessarily imply protein complex formation, hence the significant fraction of co-essential pairs with low complex quality scores. Co-essential genes with compelling associations, and yet low complex quality scores, can still be used for hypothesis generation and experimental exploration, like our itaconate defense module.

In conclusion, we provide a computational approach that relies on experimental observations to guide function assignment to orphan enzymes in *Mtb. Mtb* is a great fit for discovery of enzyme function with this approach because of the significant fraction of annotated enzymes to hint at the function of the unannotated ones. As more enzymes are annotated and more TnSeq datasets are incorporated, we expect our approach to increase in ability to identify co-essential gene clusters. We envision our approach being used throughout *Mtb* biology to speed up hypothesis generation and preliminary confirm these hypotheses. To support this vision, we have integrated a co-essentiality analysis feature into the MtbTnDB, offering an interactive platform for researchers to interactively explore gene essentiality correlations.

## Methods

### Clustering of genes using generalized least squares (GLS)

We performed unsupervised machine learning on the standardized TnSeq profiles by closely following the generalized least squares (GLS) approach of Wainberg et al. ^9^. In the GLS approach, genes are clustered based on the p-value of the GLS regression between their essentiality profiles. In our dataset, a gene’s essentiality profile is represented by its vector of differential log2 fold-changes 125 control-vs.-experimental conditions. To account for potential batch effects (i.e. non-independence of different TnSeq screens), the GLS approach incorporates the 125 ⨉ 125 covariance matrix of the differential essentially profiles - Σ - into the calculation of regression parameters. Running GLS on all pairwise combinations of genes results in a 3959 x 3959 matrix of regression p-values, which are then converted to false discovery rate (FDR) corrected q-values (qFDR) using the Benjamini-Hochberg (FDR) correction.

### Mycobacterial strains and media

*H37Rv* and its derivatives were cultured in Middlebrook 7H9 broth (Difco Laboratories) supplemented with 0.2% glycerol, 0.04% Tyloxapol and 10% ADN (50 g/L Bovine Albumin Fraction V, 20 g/L D-glucose, 8.1 g/L NaCl) in tissue culture flasks at 37° C and 5% CO_2_. For growth assays on propionate, 7H9 broth was supplemented with 0.04% tyloxapol, 0.02% propionate, 10 mg/mL of adenosylcobalamin, and 1 or 10 uM itaconate. When appropriate, antibiotics were supplemented in the following concentrations: 500 ng/mL anhydrotetracycline (ATC) and 50 ug/mL Kanamycin. For plated *Mtb*, cells were grown on Middlebrook 7H10 agar (Difco Laboratories) supplemented with 0.5% glycerol, 10% ADN, and appropriate antibiotics.

### Construction of ATC-inducible CRISPRi KD strains

For each gene targeted, CRISPRi plasmids were cloned using the PLJR965 backbone (Addgene #115163) as previously described ^69,70^. Briefly, sgRNAs were designed using Pebble’s sgRNA Design Tool to target the non-template strand of the gene ORF ^71^. (Supplementary Table 1). Two complementary oligonucleotides with appropriate overhang and sticky regions with respect to each sgRNA were annealed and ligated into a BsmBI=v2 digested PLJR965 backbone. Ligated plasmids were transformed into DH5-alpha *Escherichia coli* using heat shock transformation for plasmid replication and extraction using the Qiagen Miniprep Kit as described (REF). Cloning was confirmed using Sanger sequencing (Eton Bioscience).

Constructed CRISPRi plasmids were electroporated into *H37Rv* as previously described ^70^. Briefly, an *H37Rv* culture was grown to an OD_600_ of 0.7-.1.1, pelleted at 4000g for 10 min, and washed three times in 10% glycerol. Washed cell pellets were resuspended in 10% glycerol to a final volume that is 5% of the starting volume. For each CRISPRi plasmid, 100-200 ng of DNA was added to 50 uL of concentrated cells and pulsed using the Gene Pulser 541 X cell electroporation system (Bio-Rad #1652660) at 2500 V, 700 Ω and 25 μF. Electropated cells were recovered in antibiotic-free 7H9 for 20 hours, and plated on 7H10 agar supplemented with 50 ug/mL Kanomycin to select for successfully electroporated strains.

### Measuring Mtb itaconate-dependent growth inhibition on propionate

For each CRISPRi KD strain, an ATC supplemented and depleted culture were expanded to an OD_600_ = 0.7-1.2 for two passages in 7H9 broth to achieve complete knockdown. ATC was repleted in ATC supplemented cultures every 5-6 days. After passage two, 750 uL of cells were pelleted at 4000 rpm for 10 min, resuspended to an OD_600_ of 1 in 7H9 broth supplemented with 0.02% propionate, and incubated at in standing eppendorfs overnight at 37C and 5% CO_2_ for preadaptation to media. Pre-adapted cells were then diluted to 1:20 in 7H9 with 0.2% glycerol or 7H9 with 0.02% propionate and 1 or 10 uM itaconate to a final OD_600_ of 0.05 and final volume of 200 uL with appropriate antibiotics in a 96-well plates. Cells were incubated in 37 C at 5% CO_2_ for 23 days. Plate OD_600_ was measured every 2-3 days followed by brief agitation of cells. ATC was repleted every 6-7 days in ATC treated cells in tandem with the same volume of PBS added to untreated cells. Measurements were taken from three technical replicates for each of the three biological replicates resulting in 9 replicates. Biological replicates are defined as a unique monoclonal colony successfully transformed with a CRISPRi construct and technical replicates are defined as independent growth curves.

The SciPy statistics package was used to visualize growth curve data on the Python 3.10.0 interface and calculate average OD_600_, t statistic, p value, effect size, and 95% confidence interval of effect size^72^. The two-tailed unpaired Independent t-test to assess statistical significance. P values for relevant figures are reported in Supplementary Table 2.

### Computational Prediction and scoring of protein heterodimers

We selected candidate protein pairs as input to our virtual screen from the set of all gene pairs with a GLS p-value below 0.001 (after applying Benjamini-Hochberg false discovery rate (FDR) correction). Given computational constraints, we also imposed a threshold on the size of the candidate protein heterodimers at 800 aminoacid residues. We ran heterodimer structure predictions using ColabFold ^73^ with the following parameters: MSA mode = MMseq2 (UniRef + Environmental); number of models=2, number of recycles=3. For each protein pair, we kept the model with the highest pLDDT score for analysis with pDockQ. To generate a control set of random protein pairs, we randomly selected proteins from the *Mtb* proteome with similar size distribution as the set of GLS-derived protein pair candidates. Specifically, for each protein pair in the real test set, we generated a random protein pair with each random protein within one percent the size of the corresponding protein in the test pair. Finally, we ran the pDockQ scoring function ^16^ on the top-scoring model for each protein pair, with the distance threshold parameter set to 8 angstroms.

## Acknowledgements

We thank Ruma Banerjee, Markus Ruetz, Squire Booker, and Michael Glickman, for helpful feedback, comments, and technical advice. We thank Thomas Ioerger for his helpful feedback on co-essentiality and the TnSeqCorr database. A.J. was supported by an HHMI Hanna Gray Fellowship. S.Z.N. was supported by a Medical Scientist Training Program grant from the National Institute of General Medical Sciences of the National Institutes of Health under award number: T32GM007739 to the Weill Cornell/Rockefeller/Sloan Kettering Tri-Institutional MD-PhD Program. K.Y.R. acknowledges support by P01 (AI143575) and U19 (AI162584).

## Supplementary Tables

**Table S1.**
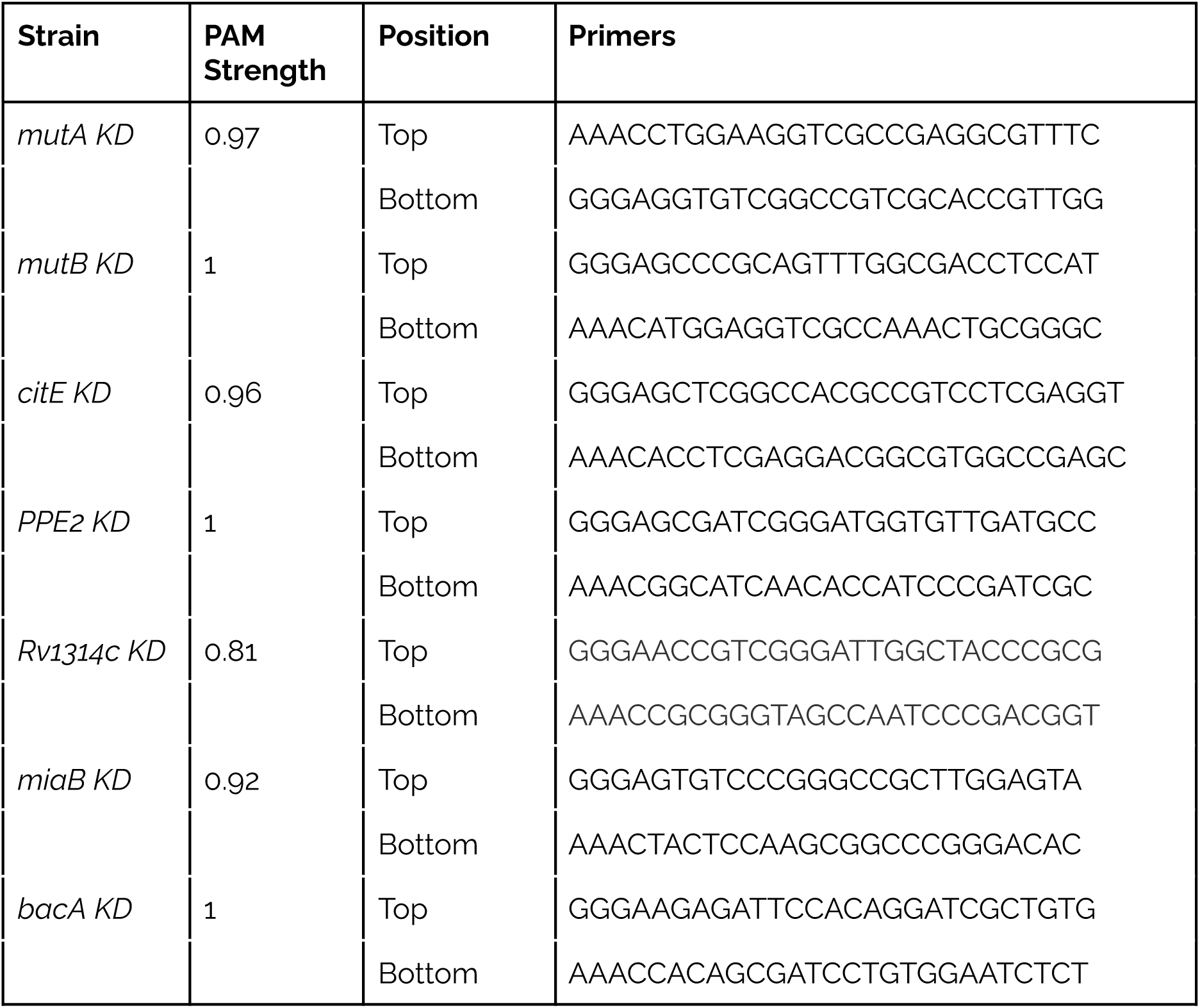
Primers used for CRISPi construction.

**Table S2.** Summary of P values for Figure 3. P values were calculated using unpaired two-sided independent t-test.

| Figure | t Test | p value | p value Summary | Degrees of Freedom | T statistic | Effect size | 95% Confidence Interval for Effect Size |
| --- | --- | --- | --- | --- | --- | --- | --- |
| Figure 3A<br>*mutA* KD | *mutA* KD +ATC vs.<br>*mutA* KD noATC | 3e-12 | **** | 8.0 | 18.68 | 0.21 | 0.18-0.23 |
| Figure 3A<br>*mutB* KD | *mutB* KD +ATC vs.<br>*mutB* KD noATC | 8e-08 | **** | 8.0 | 9.21 | 0.21 | 0.15-0.26 |
| Figure 3A *citE*<br>KD | *citE* KD +ATC vs. *citE*<br>KD noATC | 1e-08 | **** | 12.0 | 10.59 | 0.13 | 0.1-0.15 |
| Figure 3A<br>*ppe2* KD | *ppe2* KD +ATC vs.<br>*ppe2* KD noATC | 2e-07 | **** | 16.0 | 8.6 | 0.15 | 0.11-0.18 |
| Figure 3A<br>*rv1314c* | *rv1314c* KD +ATC vs.<br>*rv1314c* KD noATC | 5e-04 | *** | 10.0 | 4.37 | 0.1 | 0.05-0.15 |
| Figure 3A<br>*miaB* KD | *miaB* KD +ATC vs.<br>*miaB* KD noATC | 5e-07 | **** | 12.0 | 8.08 | 0.11 | 0.08-0.14 |
| Figure 3A<br>*bacA* KD: | *bacA* KD +ATC vs.<br>*bacA* KD noATC | 4e-08 | **** | 12.0 | 9.68 | 0.13 | 0.1-0.16 |
| Figure 3C<br>*mutA* KD | *mutA* KD +ATC vs.<br>*mutA* KD noATC | 5e-05 | **** | 15.0 | 5.62 | 0.13 | 0.08 – 0.18 |
| Figure 3C<br>*mutB* KD: | *mutB* KD +ATC vs.<br>*mutB* KD noATC | 2e-08 | **** | 15.0 | 10.1 | 0.17 | 0.14 -0.21 |
| Figure 3C *citE*<br>KD | *citE* KD +ATC vs. *citE*<br>KD noATC | 0.82 | ns | 9.0 | 0.23 | 0.01 | -0.06 – 0.08 |
| Figure 3C<br>*ppe2* KD | *ppe2* KD +ATC vs.<br>*ppe2* KD noATC | 0.07 | ns | 14.0 | 1.92 | 0.05 | -0.01 – 0.1 |
| Figure 3C<br>*rv1314c* | *rv1314c* KD +ATC vs.<br>*rv1314c* KD noATC | 4e-03 | ** | 15.0 | 3.42 | 0.09 | 0.03 – 0.14 |
| Figure 3C<br>*miaB* KD | *miaB* KD +ATC vs.<br>*miaB* KD noATC | 1e-08 | **** | 13.0 | 10.53 | 0.2 | 0.16 – 0.25 |
| Figure 3C<br>*bacA* KD | *bacA* KD +ATC vs.<br>*bacA* KD noATC | 0.82 | ns | 10.0 | -0.24 | 0.8143 | -0.01 -0.08 |
| Figure 3D<br>*mutA* | 1 μM itaconate vs.<br>glycerol | p <<br>1e-09 | **** | 11.0 | 22.52 | 1.43 | 1.29 -1.57 |
| Figure 3D<br>*mutB* | 1 μM itaconate vs.<br>glycerol | p <<br>1e-09 | **** | 8.0 | 12.25 | 1.38 | 1.12 -1.64 |
| Figure 3D *citE* | 1 μM itaconate vs.<br>glycerol | 2e-06 | **** | 12.0 | 7.3 | 0.75 | 0.53 -0.98 |
| Figure 3D<br>*ppe2* | 1 μM itaconate vs.<br>glycerol | 1e-05 | **** | 9.0 | 6.13 | 0.7 | 0.44 -0.96 |
| Figure 3D<br>*rv1314c* | 1 μM itaconate vs.<br>glycerol | 0.01 | ** | 10.0 | 2.95 | 0.4 | 0.1 -0.7 |
| Figure 3D<br>*miaB* | 1 μM itaconate vs.<br>glycerol | 9e-04 | **** | 11.0 | 4.08 | 0.37 | 0.17 -0.56 |
| Figure 3D<br>*bacA* | 1 μM itaconate vs.<br>glycerol | p <<br>1e-09 | **** | 16.0 | 13.49 | 0.9 | 0.75 -1.04 |

Values of $p \leq 0.05$ were considered significant (\* $p \leq 0.05$ ; \*\* $p \leq 0.01$ , \*\*\* $p \leq 0.001$ , \*\*\*\* $p \leq 0.0001$ ), $p > 0.05$ were considered not significant (ns).

**Table S3.** Summary of P values for Figure 3. P values were calculated using Generalized Least Squares and corrected for false discovery rate (FDR) using the Benjamini-Hochberg correction.

| Gene | Partner Gene | P value |
| --- | --- | --- |
| Rv0256c | Rv1314c | 1e-04 |
| Rv0256c | Rv1492 | 5e-04 |
| Rv0256c | Rv1493 | 2e-11 |
| Rv0256c | Rv1819c | 2e-05 |
| Rv1492 | Rv1493 | 1e-05 |
| Rv1493 | Rv1819c | 2e-05 |
| Rv1493 | Rv2733c | 9e-05 |

